# RAX2: genome-wide detection of condition-associated transcription variation

**DOI:** 10.1101/013201

**Authors:** Yuan-De Tan, Jixin Deng, Joel R. Neilson

## Abstract

Almost all of mammalian genes have mRNA variants due to alternative promoters, alternative splice sites, and alternative cleavage and polyadenylation sites. In most cases, change in transcript due to choosing alternative cleavage and polyadenylation sites does not lead to change in protein sequence, while selection of alternative promoters and alternative splice sites would alter protein sequence. Nevertheless, all these alternations would give rise to different RNA isoforms. Selection of alternative RNA isoforms has been found to be associated with change in condition. For example, many studies have revealed that alternative cleavage and polyadenylation (poly(A)) are correlated to proliferation, differentiation, and cellular transformation. Thus, unlike gene expression in microarray, change in usage of splice sites or poly(A) sites associated with conditions does not involve differential expression and hence cannot be detected by differential analysis methods but can be done by association methods. Traditional association methods such as Pearson chi-square test and Fisher Exact test are single test methods and do not work on the RNA count data derived from replicate libraries. For this reason, we here developed a large-scale association method, called ranking analysis of chi-squares (RAX2). Simulations demonstrated that RAX2 worked well for finding association of changes in usage of poly(A) sites with condition change. We applied our RAX2 to our primary T-cell transcriptomic data of over 9899 tags scattered in 3812 genes and found that 1610 (16.3%) tags were associated in transcription with immune stimulation at FDR < 0.05 and most of these tags associated with stimulation also had differential expression. Analysis of two and three tags within genes revealed that under immune stimulation, short RNA isoforms were significantly preferably used. Like cell proliferation and division, short RNA isoforms are highly prioritized to be used for cell growth.

---

Recently, alternative splicing and alternative cleavage and polyadenylation (ACP) have been revealed to be not only a universal posttranscription processing step in eukaryotic gene expression but also a versatile mechanism for posttranscriptional regulation of genes (Colgan and Manley 1997; Shepard et al. 2011; Zhao et al. 1999). After transcription, a pre-mRNA is capped at 5’ end, spliced, and cleaved in the 3'-untranslated region (3'UTR) to yield a new open end that allows to add a polyadenylation (poly(A)) tail(Shen et al. 2011; Zhao et al. 1999). A poly(A) tail at 3'end may protect the mRNA from unregulated degradation, trigger export of the mRNA to cytoplasm, and assist recognition by translation machinery(Danckwardt et al. 2008; Shen et al. 2011; Zhao et al. 1999). A poly(A) signal at which the pre-mRNA is cleaved and referred to as poly(A) site is recognized and activated by a balk of protein factors called polyadenylation factors(Hunt et al. 2008; Shen et al. 2011; Shi et al. 2009; Zhao et al. 1999). Alternative splice and poly(A) sites significantly increase complexity of transcriptomes and proteomes because they lead to multiple isoforms or variants of a mRNA. Using next generation sequencing (NGS) techniques, it has been observed that over 50% of the transcriptome in the mammalian genome have alternative cleavage and polyadenylation (Ji et al. 2009; Lee et al. 2007; Meyers et al. 2004; Shen et al. 2008; Tian et al. 2007; Wu et al. 2011). As alternative splice and poly(A) sites can uncover variation of transcription at subgene level, more and more investigators attempted to find transcription variation associated with tissue types or cell states and explore etiologic mechanism of disease of interest in transcription variation (Flavell et al. 2008; Ji et al. 2009; MacDonald and McMahon 2011; Mayr and Bartel 2009; Sandberg et al. 2008; Wang et al. 2008; Winter et al. 2007; Yu et al. 2006). But almost all these studies are based on differential analysis. This may be because the current large-scale statistical methods such as DESeq(Anders and Huber 2010), baySeq(Hardcastle and Kelly 2010), edgeR exact test (Robinson et al. 2009; Robinson and Smyth 2008), edgeR GLM (McCarthy et al. 2012), DEXSeq (Anders et al. 2012), Cuffdiff (Trapnell et al. 2013; Trapnell et al. 2010), DiffSplice(Hu et al. 2012), SplicingCompass (Aschoff et al. 2013) are differential analysis methods. Of these methods, the former fours are used to identify differential transcription between conditions and the latter fours are used to find differential splicing between cconditions. However, unlike differential expression of genes between conditions, splice switch of alternative splice sites or usage switch of alternative poly(A) sites occurs on the same transcript unit (Fig.1). So to use the differential analysis methods to identify usage and splicing switches from site to site on the RNA unit is not a good way. While association analysis approaches are useful to address this kind of problem, to our best knowledge, no existing large-scale association analysis methodologies have been proposed. The traditional association methods such as chi-square test and Fisher exact test are single-test methods and works on the count data without replicates, they hence could not directly be applied to high-throughput transcriptomic data derived from replicate libraries. This is why we need to develop a large-scale chi-square statistical method for identifying tissue-associated or condition-associated isoform transcriptions. This method is named ranking analysis of chi-squares (RAX2). RAX2, which extends traditional chi-square test to replicated and high-dimensionality transcriptomic data, is based on comparison between a set of ranked chi-square statistics for association effects and a set of ranked null chi-square values across a set of given thresholds and estimation of a false discovery rate (FDR) profile.

**Figure 1.**
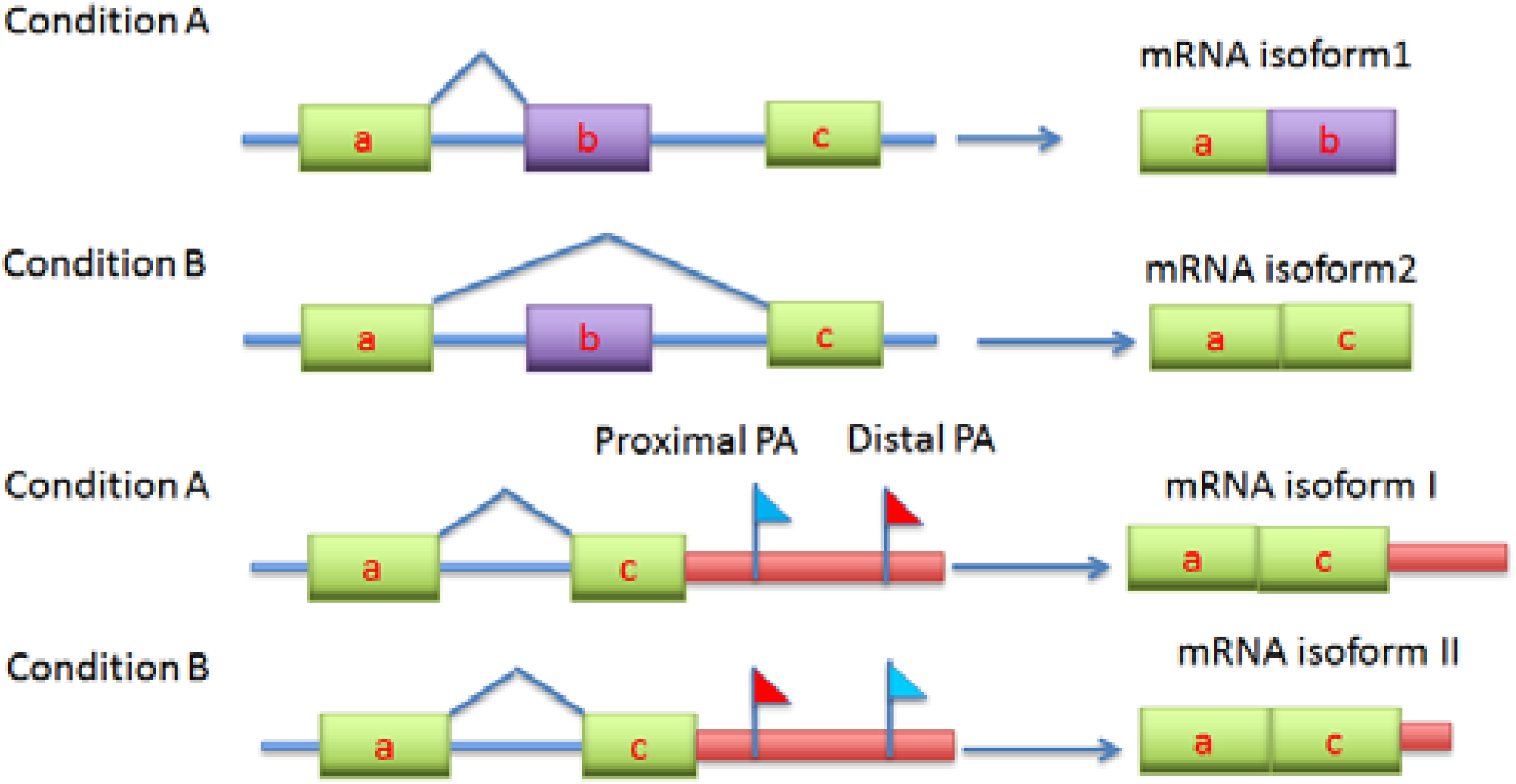
Demonstration for association of usage of alternative splice and poly(A) sites with conditions: splicing switch and Usage switch of poly(A) sites due to change in condition. Under normal condition, the gene transcription product is isoform 1 due to splicing event between exon a and b, but in stress or stimulation, for example, the gene product is isoform 2 by switching splicing from exon b to exon c. This is called splicing switch. Likewise, under normal condition, cells use distal poly(A) site and the transcription produces mRNA isoform I but when cells are stressed or stimulated, poly(A) site usage switches from distal site to proximal site and produces isoform II. This is called poly(A) site usage switch.

## Results

### Models

For the convenience, our discussion about RAX2 is based on alternative cleavage and polyadenylation (ACP) sites within genes. ACP can occur concomitantly with or independently of alternative splicing (Fig.2). ACP independent of alternative splicing, that is, all cleavage and polyadenylation (poly(A)) sites are in the terminal exon of the transcription unit, results in a transcription unit with a tandem untranslated region (tandem UTR). Both alternative splicing of terminal exons and tandem UTR usage are visible for directed 3' end sequencing methodologies. Since either process has the potential to alter 3' UTR identity, we do not differentiate between them in this analysis. For a given transcription unit, transcript variants derived from the first poly(A) site (poly(A) site 1) are assumed to have a transcript length from the transcriptional start position to poly(A) site 1. Similarly, transcript variants with poly(A) sites 2 and 3 are also assumed to be derived from the transcriptional start position to poly(A) sites 2 and 3, respectively. Therefore, within such a transcription unit, the transcript isoforms are one-to-one corresponding to poly(A) sites. For the sake of convenience, we refer to the transcript variant at poly(A) site 1 as tag1. Similarly, the transcript variants at poly(A) sites 2 and 3 are also defined as tags 2 and 3, respectively (Fig.3). In general, within gene g, *Z_g_* alternative poly(A) sites correspondingly have *Z*_*g*_ alternative tags. For example, gene AARS2 on chromosome 6 has 4 poly(A) sites(*Z*_*g*_=4) and hence its mRNA transcription unit (called gene) has 4 isoforms (see columns gene name and position(pos) in Supplemental Table S1). Table S1 lists such count data of reads of tags within genes where columns 1-7 list information of tags or poly(A) sites within genes: tagid, geneid, name, DNA strand, position on transcription unit, and annotation; columns 8-10 list count data of cells in normal state and stimulation. The data were normalized so that they look decimal. Here we let *n*_*gzjv*_ be count of tag *z* (*z* = 1,…, *Z*_*g*_ ≥ 2) within gene *g* (*g* = 1, 2,… *G*) in cell state *j* in replicate *v* (*v* =1,…, *r*). Our concern here is if change in count of a tag *z* at a special poly(A) site *within* a gene of interest is associated with change in condition (cell state) in the context of a discontinuous process. For the convenience, we focus on two states: a treatment state (TS) and a normal state (NS). We set *j*=1 for NS and *j*=2 for TS. To find if the transcription variation of an individual tag is associated with a change in cell state, we need to compare count of tag *z* to that of all the other tags (denoted by 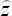) within gene g. Therefore, a tag or poly(A) site has two states: z and 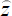. We set *i*=1 for z and *i*=2 for 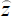). Thus count *n*_*gzjv*_ in the raw data table can be converted into two-by-two table count *n*_*gzijv*_ where *n*_*gzijv*_ is count of reads of tag *z* within gene *g* in tag state *i* in condition *j* in replicate *v* (*v* =1, 2, …, *r*) and there are *Z*_*g*_*r* two-by-two table datasets (*n*_*gz*11*v*_, *n*_*gz*12*v*_, *n*_*gz*21*v*_, *n*_*gz*22*v*_) for gene *g*. Since our interest is in regards to tag *z* (or poly(A) site *z*) within gene *g* instead of gene g itself, totally we have 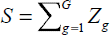 tags or poly(A) sites in all genes of study. To implement ranking analysis of all tags, we combine subscripts *g* and *z* into *s* where *s* =1, …, S. Thus, each tag *s* has two states (s and 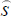). We still set *i*=1 for tag state s and *i*=2 for tag state 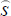. The two-by-two table dataset is then rewritten as (*n*_*s*11*v*_, *n*_*s*12*v*_, *n*_*s*21*v*_, *n*_*s*22*v*_).

**Figure 2.**
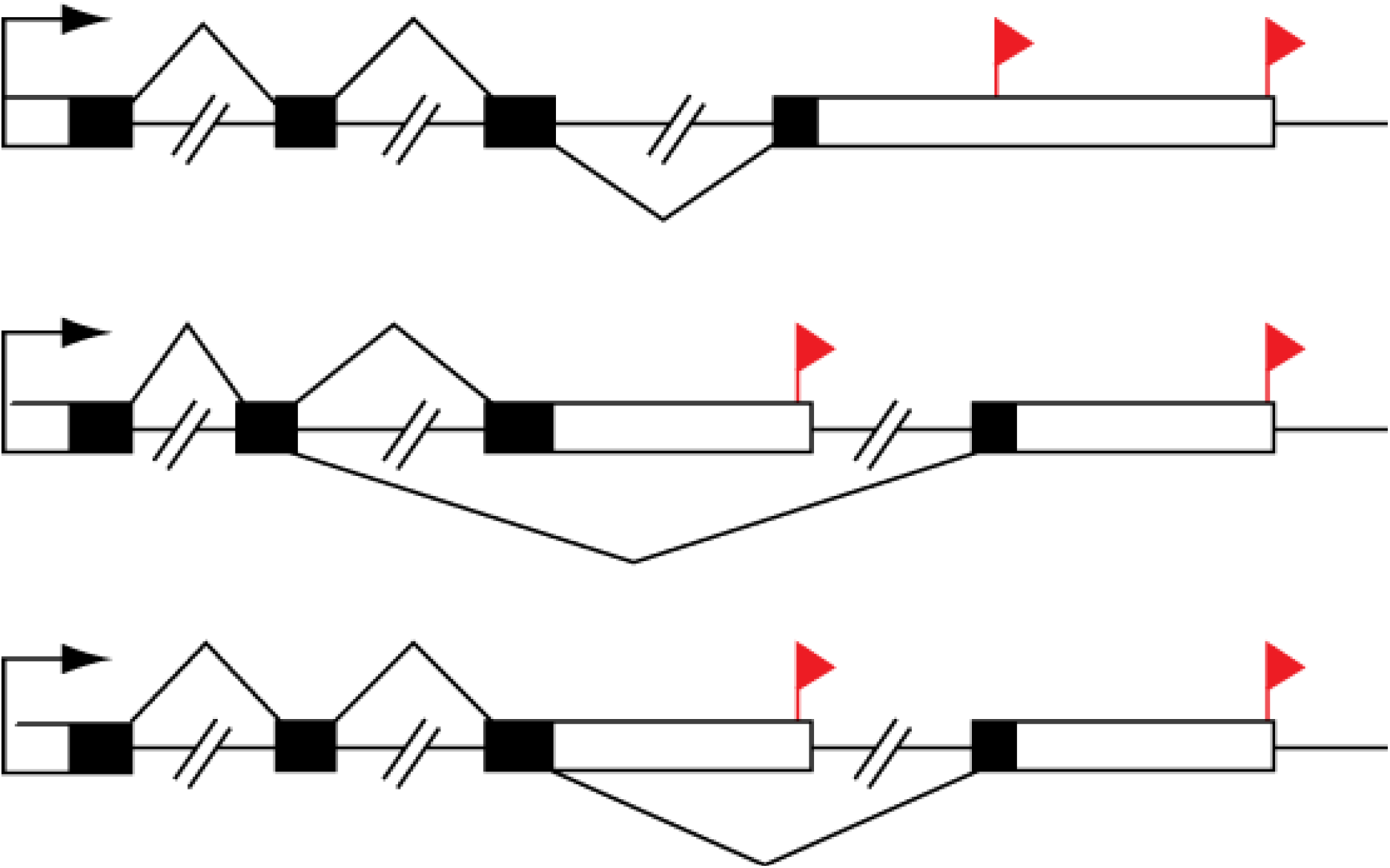
Transcription unit structures and different 3' UTR isoforms. A majority of mammalian transcription units are characterized by alternative cleavage and polyadenylation (ACP). The majority of these contain multiple cleavage and polyadenylation sites in their terminal exon (top), impacting untranslated region identity without changing the coding sequence. Transcription units may also be characterized by mutually exclusive terminal exon structure (middle) or composite terminal exon structure (bottom). In the latter two cases, ACP is coupled to mRNA splicing, and both the coding sequence and the untranslated region are impacted. Black box: exon, white box: untranslated region(UTR), red flag: poly(A) site and -//-: intron.

**Figure 3.**
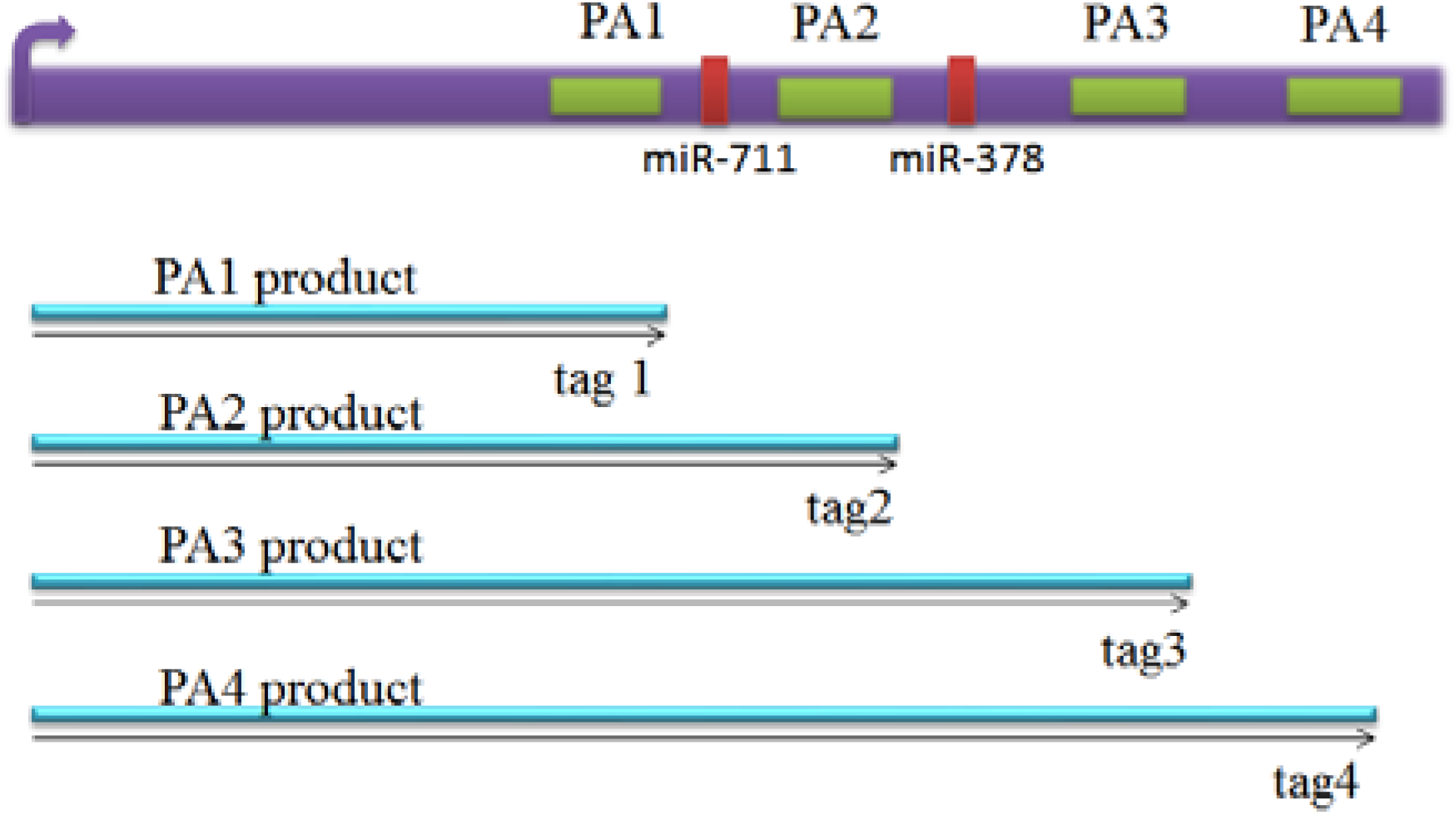
A model for multiple poly(A) sites or tags within genes. mRNA of Hsp70.3 is a typical model for multiple poly(A) sites (green boxes) and microRNA sites (red boxes). It has a common start point (promoter 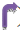), four poly(A) sites (PA1, PA2, PA3 and PA4) in 3’UTR coding for several mRNA isoforms that are different in their 3'end. Alternative polyadenylation changes the length of the 3'UTR and it also can change microRNA binding sites in 3'UTR. While microRNAs tend to repress translation and promote degradation of the mRNAs they bind to. Since transcription products are one-to-one corresponding to poly(A) sites, we define the transcription products as tags.

Since sample sizes for each two-by-two table are small, the observed values consist of two components: the association effect and random noise. Thus a model for each observed value in such a two-by-two table is

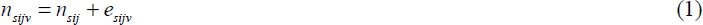

where *n*_*sij*_ is count of reads of tag *s* due to association effect between state *i* of poly(A) site *s* and cell state *j* and *e*_*sijv*_ is count of reads due to a special noise between state *i* of poly(A) site *s* and cell state *j* in replicate observation *v*. Since *n*_*sijv*_ follows Poisson distribution (Anders and Huber 2010; Robinson and Smyth 2008) or negative binomial distribution (Anders and Huber 2010; McCarthy et al. 2012; Robinson and Smyth 2008) or binomial distribution(Baggerly et al. 2003), *e*_*sijv*_ may also follow negative binomial or binomial distribution. But we do not concern distribution of *e*_*sijv*_ because we do not need to separate it from *n*_*sijv*_. Let *f*_*sji*_ and *f*_*sijv*_ be frequencies of counts *n*_*sij*_ and *e*_*sijv*_, respectively. As *n*_*sij*_ is count of reads of tag *s* in state *i* and in cell state *j*, *n*_*sij*_ = *n*_*s*_*f*_*sij*_ and *e*_*sijv*_ = *n*_*s*_*f*_*sijv*_ where *n*_*s*_ is total count of reads of tag *s* within *g* across two cell states and two tag states over all replicate libraries. Thus, model (1) may be rewritten as

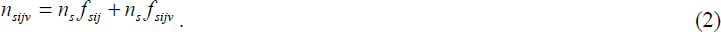

### Chi-square statistics

For poly(A) site *s*, the count of reads due to association effect *n*_*sij*_ between site state *i* and cell state *j* is estimated by

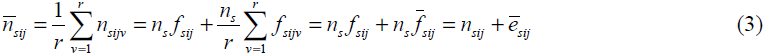

where *f*_*sij*_ is the frequency of association effect *n*_*sij*_ between state *i* of tag *s* and cell state *j* and *f*_*sijv*_, frequency of a special noise in replicate observation *v*. For poly(A) site *s*, 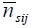 is expected as

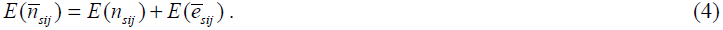

(See Supplemental Note S1 for detail). Thus, Pearson chi-square statistics for mean count 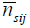 that is estimate of association effect *n*_*sij*_ is

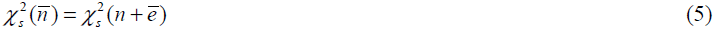

(See Supplemental Note S2 for derivation). From Equation 5, we can see that 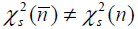 unless the mean of noises is zero 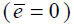, meaning that the Pearson chi-square for 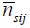 is not an unbiased estimate of the chi-square statistic for *n*_*sij*_. For this reason, we cannot use 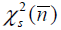 as standard chi-square statistic, that is, p-value for 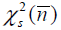 obtained from a standard chi-square distribution has a big bias. In addition, 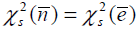 if, for a tag *s*, there is no association effect *n_sij_* between tag states and cell states, say, 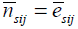.

### Null chi-squares

Similarly to using within-group variance to estimate null variance, we also employ within-group chi-square to estimate null chi-square. To this end, we need to construct another contingent table in which we have site states *s* and 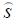) within gene *g*, and *r* replicates in cell state *j*. Set *i*=1 for s and *i*=2 for 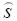. A set of two-by-two tables (see Supplemental Note S3) is constructed with pairs of replicates and states of tag *s* in cell state *j*. Thus the null chi-square is easily estimated by 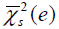 (see Supplemental Note S3 for derivation).

### Ranking analysis of chi-squares

Since 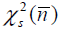 does not have one-to-one correspondence to 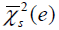, to compare them, we need to separately rank tags by amounts of 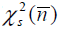 and 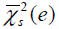 from the smallest to the largest. Let 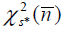 and 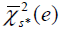 be two Pearson chi-square values at position *s*^*^ in the ranking space(*), the chi-square values are smallest at *s*\*=1 and largest at *s*\*= *S\**. Then we can compare 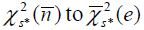 at position *s*\* in a one-to-one fashion. Given thresholdΔ, be associated with change in cell state if and only if, expression change of tag *s\** is declared to be associated with change in cell state if and only if

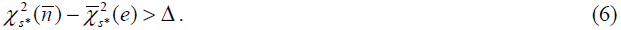

### FDR estimation

In large-scale statistical analysis, we do not need to concern about so-called type 1 error but we must consider how to control false discovery rate (FDR) because in, for example, 10000 hypotheses to be tested, at least 500 hypotheses would commit type I error if α=0.05 is chosen as significance level(Tusher et al. 2001). Obviously 500 hypotheses rejected by chance are not acceptable. This is why we here want to control FDR rather than type I error. A reasonable control of FDR involves a reliable estimation of FDR. Given threshold Δ, the number (*N*_Δ_) of tags whose expression changes are declared to be uniquely associated with change in cell state may consist of the number of truly positive tags, *T*_Δ_, and number of falsely positive tags, *F*_Δ_:

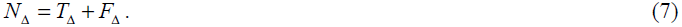

Therefore, FDR is expected as *FDR*_Δ_ = *E*(*F*_Δ_ / *N*_Δ_) at threshold Δ. As *F*_Δ_ is unknown in experimental data sets, FDR must be estimated.

The existing methods for estimation of FDR are not suitable to our current chi-square statistics because, as seen above, chi-square statistics are remarkably biased against standard chi-square statistics such that p-values obtained from standard chi-square distribution are also biased. For this reason, we here propose a novel method. The principle of the method is shown in Figure 4. As seen in Figure 4, observed chi-square 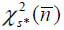 more and more deviates from expected chi-square 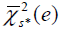 as chi-square value increases.

**Figure 4:**
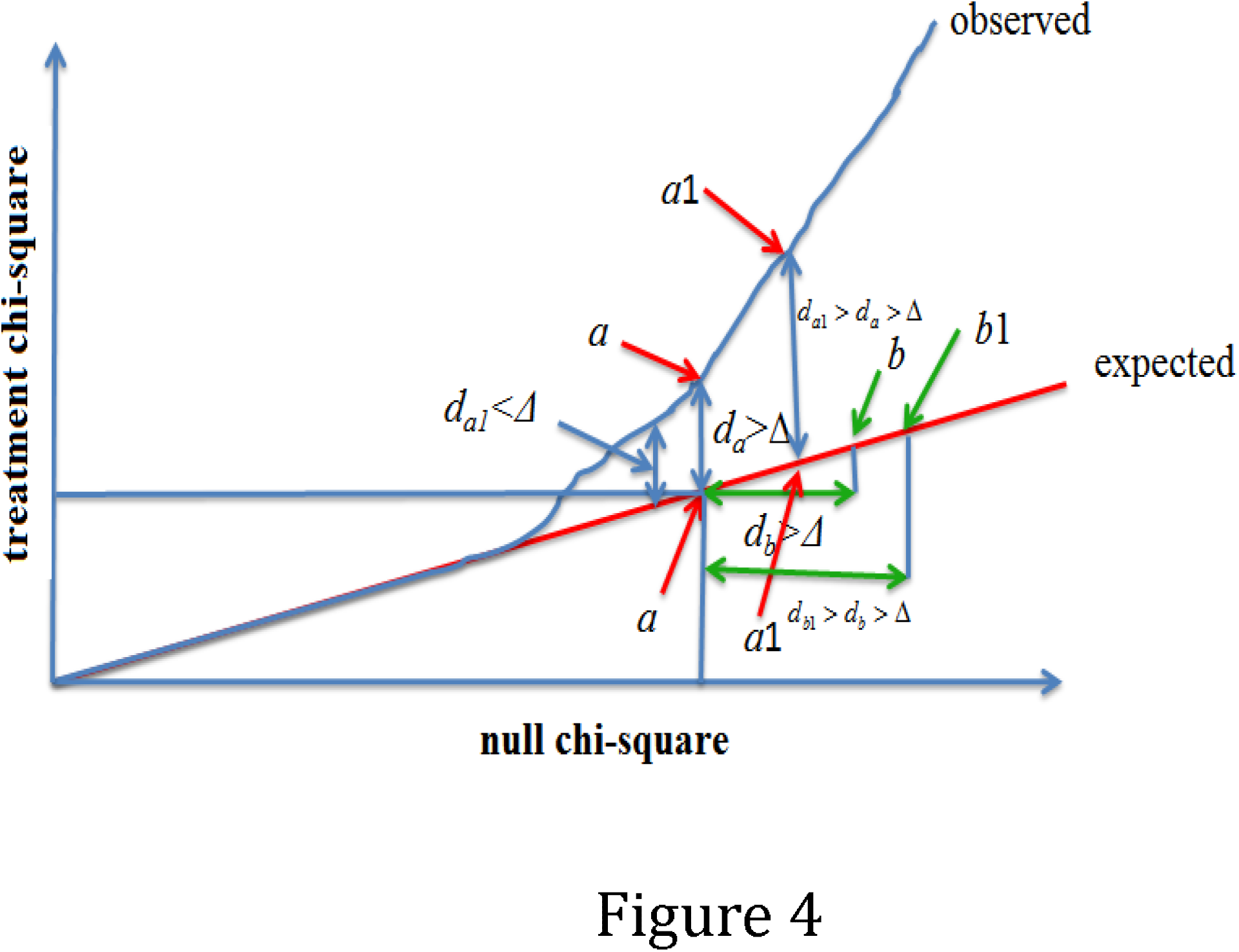
A demonstration plot of ranked treated chi-square versus null chi-square The expected linear plot (red line) is given by null hypotheses that the treated chi-square is equal to null chi-square at each chi-square point. The observed linear plot is given by ranked observed treated chi-squares versus ranked estimated null chi-squares. Given a threshold Δ, all treated chi-squares with *d*_*s*_ ≥ *d*_*a*_ are declared to be significant or interested where 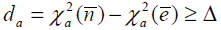 and s =a+1,…, G. All chi-squares with *d*_*t*_ ≥ *d*_*b*_ are defined as potential false positives with probability given in Equation 12 where 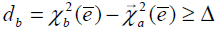and t=b+1,…, G.

Given threshold Δ, we find that 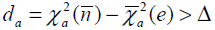at *s*\* = *a* is smallest among all *s*^*^. We define *d*_*a*_ as *d*_Δ_ and *a* as *a*_Δ_. All 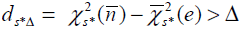 would be larger than or equal to *d*_Δ_ where *s*\* = *a*_Δ_, *a*_Δ_ + 1, .., *S*, that is, all *s*\* ≥ *a*_Δ_ would be declared as positive tags. So, the number of positives declared by Equation 6 is obtained by:

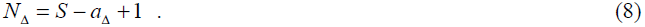

Likewise, for threshold Δ, at *t*\* = *b*_Δ_, we also find that 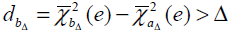 is smallest among all *t*^*^. All 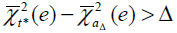 would be larger than or equal to *d*_*b*_Δ__. In addition, within the ranking space of 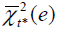, null tag at *t*\* = *b*_Δ_ + *j* has smaller chance to be chosen as a false positive than at *t*\* = *b*_Δ_ + *k* where j < *k* = 1, … *S* − *b*_Δ_ and the ranking space of 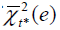 varies with sample size. In other words, the probability that a null tag at position *t*\* is chosen as false positive is determined by replicate number *r*, chi-square space, tag space *S\**, null chi-square values, and *t*\*. The logic relationship is that position *t*\*in the ranking space is positively related to the probability that the null tag at position *t*\*is chosen as a false positive tag and null chi-square value is also positively related to the probability that the null chi-square is chosen as false positive chi-square. As position *t*\* is fixed in a given tag space, the ratio ( *t* * / *S* *) would not fitvto changeable false positive probability from practical data; while 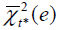 is variable and dependent on a practical dataset, 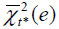 at some nearby positions may not vary or very approximate, so ratio 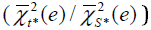 is also not appropriately used as the false positive probability. To obtain an accurate false positive probability we need to combine them. A good and simple combination way is geometric mean: 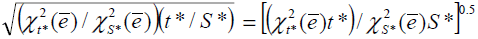. Additionally, in single hypothesis test p-value of statistic is negatively related to sample size, similarly, the probability is also negatively related to sample size because sample size increment would let space of 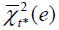 decrease. On the other hand, 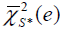 is positively related to the probability. We therefore replace 0.5 with 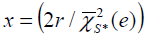. Thus, probability that tag at position *t* * is chosen as a false positive tag is given by

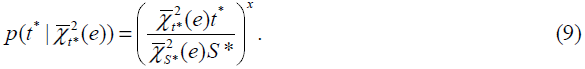

Number of false positives in *N*_Δ_ findings is estimated by

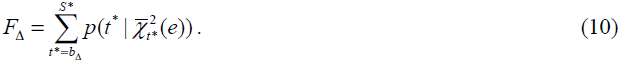

When *x* → 0, 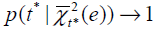 such that *F*_Δ_ → *S* − *b*_Δ_ + 1. This is an extreme case in which FDR is very conservatively estimated (see Supplemental Fig.S1A1, A2, and A3). In general, larger sample sizes would have a smaller 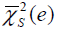, leading to a larger *x* value that causes a small 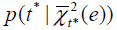. This is agreeable with the fact that larger sample sizes would have a smaller FDR. With *F*_Δ_, FDR for *N*_Δ_ findings declared by chi-square tests at threshold Δ is estimated as

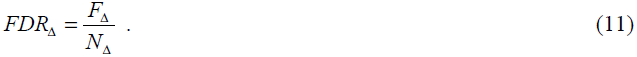

### Simulation evaluation of RAX2

We use the following steps to generate the null data of *S* tags with *r* replicates based on the real data in a group:

Step1: Calculate variance and mean of each cell in two-by-two tables over *r* replicates.

Step2: Choose randomly one of four variances and one of four means.

Step 3: Generate a set of two-by-two random null count data (*η*_*sijv*_) with *r* replicates using negative binomial pseudorandom generator in R with the mean as mu and the variance as dispersion (size).

Step 4: Repeat steps 1-3 until *s* = *S*.

We then adopted uniform and normal pseudorandom generators to generate count data with association effects:

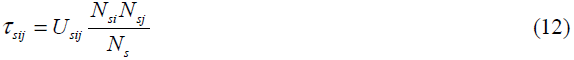

where *N*_*si*_ and *N*_*sj*_ are normal variables with *σ* =50 and mean =100 (these values are arbitrary because associate effect does not depend on individual values), 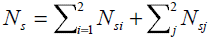, and *U_sij_* = *U_si_U_sj_* where *U_si_* and *U_sj_* are uniform variables, 0 < *U_si_* ≤ 1 and 0 < *U_sj_* ≤ 1, *i*=1 and 2, *j*=1 and 2, *s* =1, …, S. The association effect *n*_*sij*_ is randomly assigned to 10%, 20%, and 30% of null tags (the null data above). Using these simulated datasets, we compared the estimated FDR given by a statistical method to its true FDR across a set of given thresholds (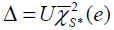, 0 ≤ *U* ≤ 1).

Supplemental Figure S1 summarizes these results in the simulated scenarios where 10% (A1 and B1), 20%, (A2 and B2), and 30% (A3 and B3) of tags have association effects on transcription with cell states and *r* = 6, *x* = 0 and 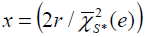 With *x* = 0 (Fig. S1A1-3), FDR was significantly overestimated in all three given scenarios such that many tags with true association effects would be missed at FDR <0.05 level. In contrast, with 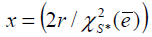 (Fig. S1B1-3), the estimated FDR curve is slightly higher than the true one in each of these three scenarios, finding tags with a true association effect at a given threshold with higher power. Supplemental Figures S1C1-3 are obtained with 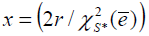 from the simulated data with *r* = 3 and 10%, 20% and 30% of tags having association effect, respectively. The fact that these FDR curves are very similar to those in Supplemental Figure S1B1, B2, and B3 indicates that our estimate of FDR with 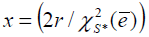 is absolutely conservative but close to its true value across a set of given thresholds in these scenarios.

### Comparison with Pearson chi-square, Fisher exact test and Cochran Mantel Haenszel (CMH) chi-square test approaches

In order to show strong advantages of our methods, we applied Fisher exact test, Pearson chi-square test and Cochran Mantel Haenszel (CMH) chi-square test approaches to the simulated data and used Benjamini-Hochberg (BH) multiple-testing procedure(Benjamini and Hochberg 1995; Benjamini and Yekutieli 2001) and q-value (Dabney and Storey 2006; Storey and Tibshirani 2003) to estimate FDR. Since the Fisher exact test and Pearson chi-square methods do not work on the replicated data, we utilized mean counts over *r* replicate libraries to construct two-by-two table for each tag. These association test methods were performed in R packages (fisher.test, chsq.test and CMH.test), the BH procedure was conducted in R package p.adjust with method = BH. Package qvalue was downloaded from Bioconductor and its qvalue.gui() was used to calculate q-values. The p-values were obtained from application of these three approaches to our simulated data of 15391 tags with 3 replicates among which 10% of tags were given with association effect values. The three sets of 15391 p-values for Fisher exact tests, Pearson chi-square tests and CMH chi-square tests were listed in Supplemental Table S2 and adjusted by HB-procedure and q-value method. Supplemental Figure S2 displays the results of plotting estimated against true FDR values across all cutoff points. One can find from Figure S2 that FDRs are completely overestimated by BH-procedure and q-value approach. The same case was also found in the simulated data with 5 replicates (data not shown). As indicated above, Pearson chi-square based on mean counts over replicates is not unbiased and the p-value for such chi-square obtained from a standard chi-square distribution is greatly biased against its true value (See Supplemental Table S2). So BH and qvalue procedures are not available to adjust such biased p-value profiles.

### RAX2 analysis of the primary T-cell data

We then assessed the performance of RAX2 on 3p-seq datasets derived from resting and stimulated primary human CD4^+^ T cells (Supplemental Table S1, see Data Collection). After normalizing and filtering, our dataset contained 16247 tags scattered in 10160 transcription units or genes. We omitted transcription units with a single tag and the remaindered 9899 tags scattered in 3812 genes were available for RAX2 analysis. We used classical chi-square distribution to calculate p-value for each tag chosen and applied the BH- procedure to adjust α-values. By comparing the ordered p-values to the ordered adjusted alpha values, as expected in Supplemental Figures S2, 747 tags were found to be associated with cell states at FDR<0.05 (see Supplemental Table S4). We performed RAX2 on our primary T-cell transcriptomic data. The estimated null chi-square value falls in a range of 0 to 6.7 (Fig. 5A). The treatment chi-square falls in interval between 0 and 363, which is larger than that in the estimated null chi-square distribution. The observed linear dots are deviated from the expected linear dots when the null chi-square is larger than 2.5 (Fig. 5B).

**Figure 5.**
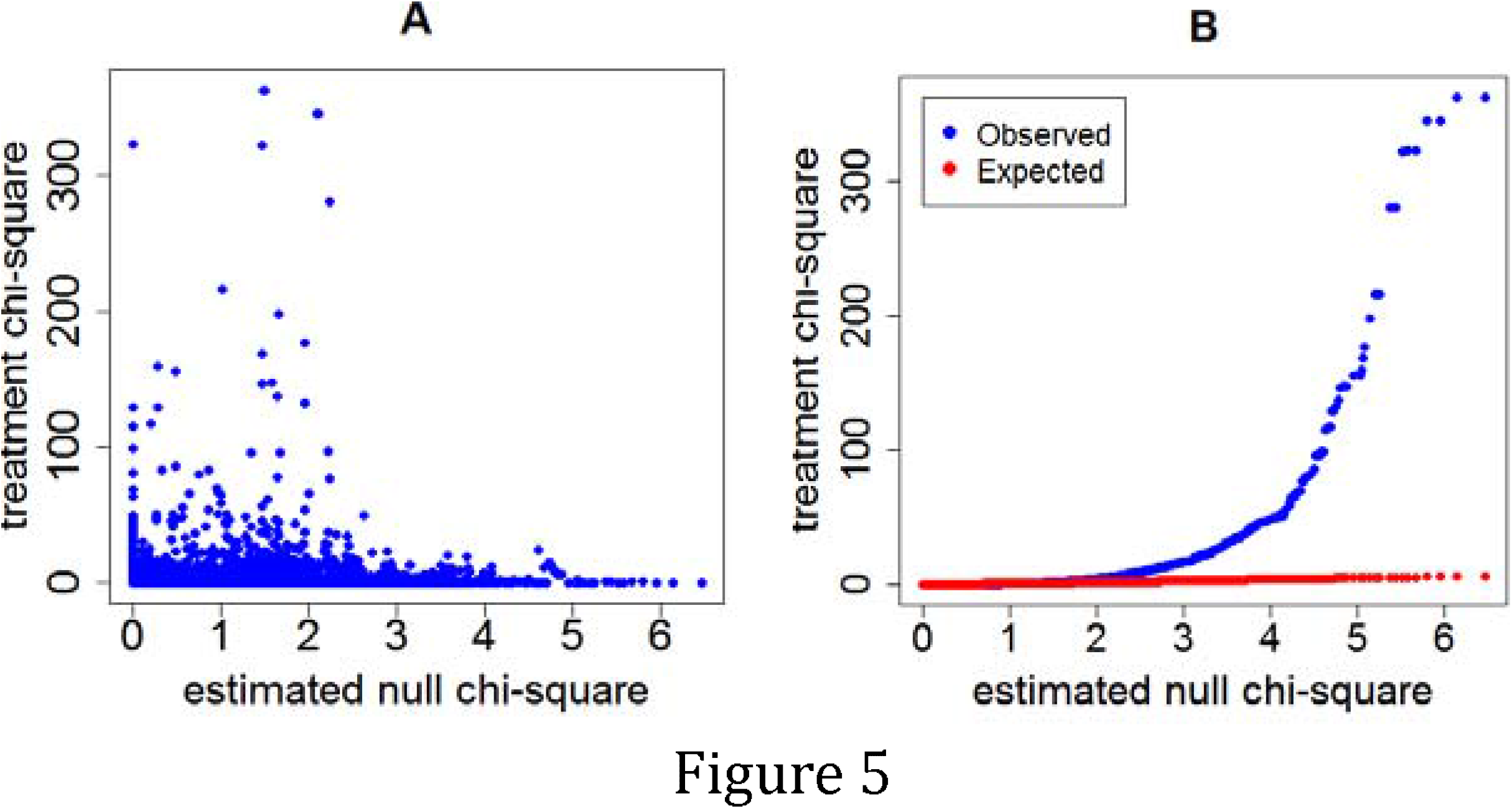
Scatter and linear plots of treated versus estimated null chi-squares. **A:** Scatter plot of treated chi-squares against estimated null chi-squares derived from primary experimental data. Over 80% of treated chi-squares fall in the null chi-square distribution. **B:** Linear plot of ranked treated chi-squares against ranked estimated null chi-squares. The observed linear dots (blue line) were significantly deviated from the expected linear dots (red line) across the estimated null chi-square distribution.

The results obtained by application of RAX2 to our real transcriptomic data sets are illustrated in Supplemental Tables S3-S5. At FDR < 0.05, 1610 (16.3%) tags were found to be associated in transcription with antigen receptor stimulation (Supplemental Table S3). Among 1610 tags, 1228 tags have fold change >1.5, meaning that most of tags found by RAX2 also displayed differential expression between rest and stimulation. We chose genes with two tags identified at FDR<0.05 (see Supplemental Table S6), and plotted logarithms of the ratio of transcription relative frequencies of proximal tag in stimulation to those in normal condition against those of distal tags and observed four patterns of transcription variation: forward and backward “switch” changes; positively and negatively accordant changes (Fig. 6A) where forward switch is defined when high expression is switched from proximal site to distal site and backward switch when high expression is switched from distal site back to proximal site. We call it positively accordant change when expression of two tags is highly raised by stimulation or negatively accordant change when both come down due to stimulation. In our data, genes with forward switch were less than genes with backward switch (148 vs 186) and positively accordant genes were many more than negatively accordant genes (186 vs 85) (Fig.6B). This means that stimulation significantly promoted transcription of tags. Proximal tags positively associated with stimulation are more than distal tags positively associated with stimulation (372 vs 334) (Fig. 6C). This result indicates that proximal poly(A) sites were preferably used under stimulation. This is consistent with the previous studies (Wang et al. 2008; Winter et al. 2007; Yu et al. 2006). Sandberg et al (Sandberg et al. 2008) found that tandem UTR length is highly negatively correlated to proliferation ratio. That is, proliferative cells more use proximal poly(A) sites. Mayr and Bartel (Mayr and Bartel 2009)also observed that cancer cells preferably used short 3’UTR while tissue cells more used long 3UTR because cancer cells have high proliferation ratio but tissue cells are highly differentiated. CD3/CD28 co-stimulation trigged a series of physiological activations and expansion (growth) of T-cells. Rapid growth and proliferation of cells require a large number of mRNAs, short mRNA isoforms can more easily meet this requirement than long isoforms, and so short mRNA isoforms are preferably produced. Figure 7 displays examples for forward and backward switches, positively and negatively accordant changes. Figure 7A is backward switch between two tags within gene PHF6 (PHD finger protein 6) potentially playing role in transcription regulation. Stimulation made high transcription level switched from distal poly(A) site backward to proximal poly(A) site. In gene SRP68 (Signal Recognition Particle of 68kDa, a ribonucleoprotein complex), the transcription pattern of two tags (Fig.7B) is completely inverse with that in gene PHF6, called forward switch. Figures 7C and D show negatively and positively accordant transcription of proximal and distal tags in genes ARL4C (ADP-Ribosylation Factor-Like 4C) and IL24 (interleukin 24). Stimulation triggered high transcription of two tags in gene IL24 but strongly suppressed transcription of proximal and distal tags in gene ARL4C.

**Figure 6:**
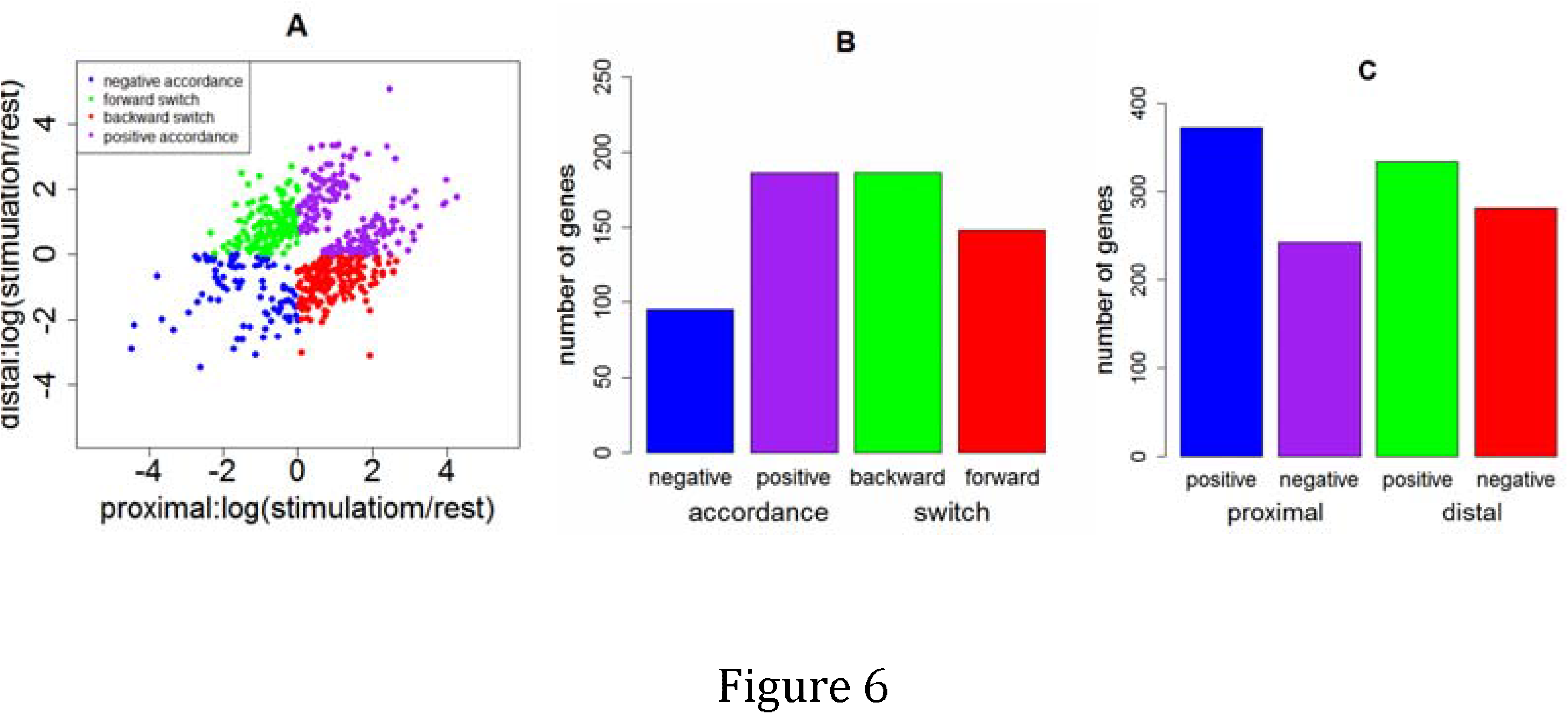
Two-way scatter plot for four association patterns of transcription of two tags within genes between two cell states. A two-way scatter plot displays distributions of scatter dots in four phases: Phase II for forward switch, phase V for backward switch, phase I for positive accordance and phase III for negative accordance. A: Plot of proximal tags versus distal tags where coordinates *x* and *y* are differences between ratios of counts of tags in stimulation and those in rest state. The ratio = sum of transcription counts of a tag over all replicates in a cell state / sum of transcription counts of this tag over all replicates and all cell states. B: Numbers of genes with four transcription patterns of proximal and distal tags. C: Numbers of genes with proximal tags positively and negatively responses to stimulation and numbers of genes with distal tags positively and negatively response to stimulation

**Figure 7.**
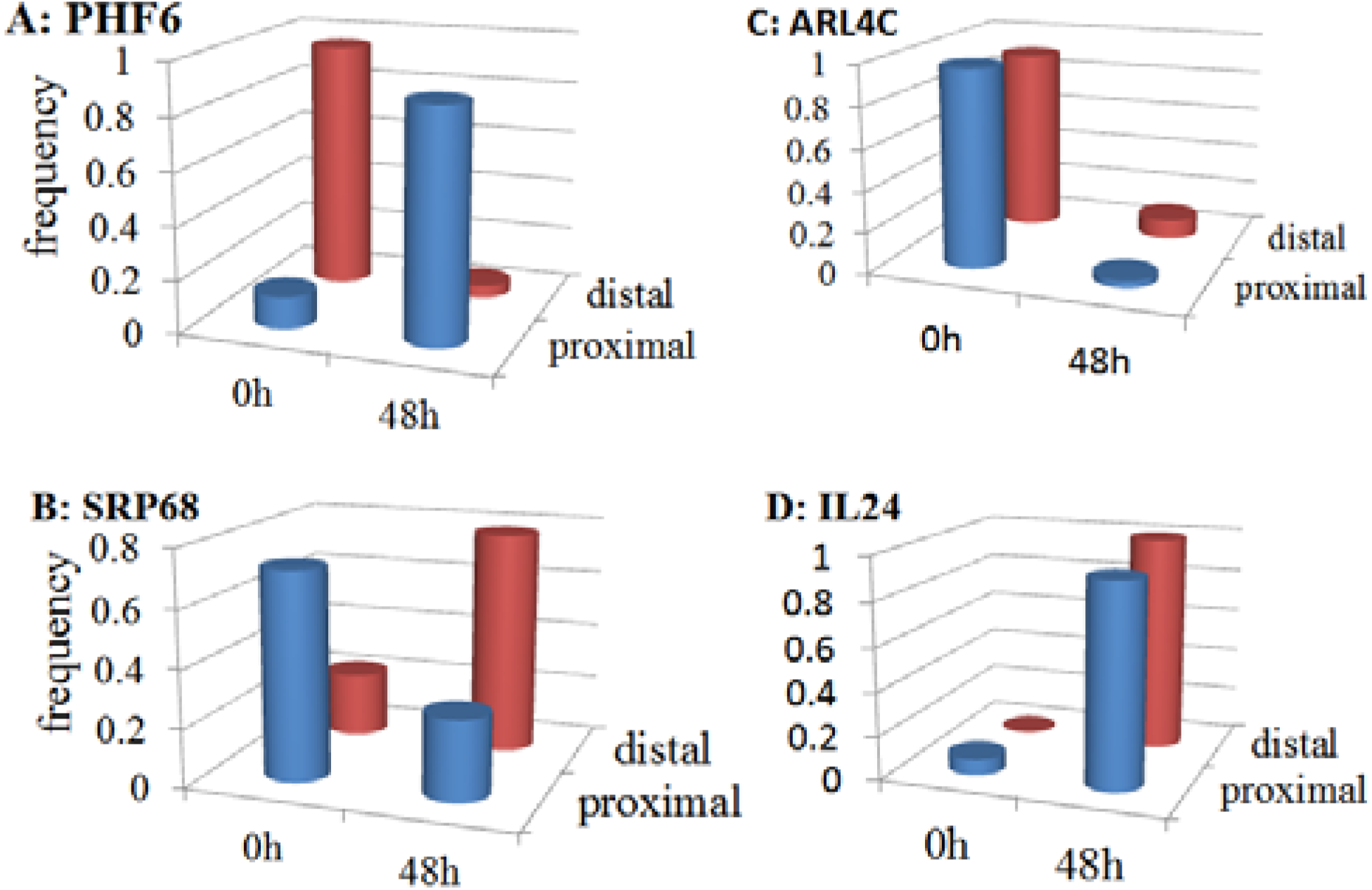
Examples for association patterns of transcriptional representation of two tags within genes between two cell states. Gene products shown here contain two tags defined by usage of distal and proximal poly(A) sites. Association between relative expression of two tags within genes and the cell states, detected by RAX2, shows backward switch (A), forward switch (B), negative accordant changes (C) and positive accordant (D). y- axis corresponds to ratio of count sum of a tag over all replicate libraries in a cell state (0 h or 48 h poststimulation) to the sum of the tag over all replicate libraries across all cell states.

In our result, 162 genes were found to have three poly(A) sites whose usage were associated with stimulation (Supplemental Table S8). Figure S3 displays three scatter plots of these genes with proximal tags versus middle tags (Supplemental Fig.3SA1), proximal tags versus distal tags (Supplemental Fig.3SA2) and middle tags versus distal tags(Supplemental Fig.3SA3). In plot of proximal versus middle tags (Supplemental Fig.3SA1), accordantly changed tags were many more than switched tags (47 vs 9 in Supplemental Fig.3SB1). Among accordantly changed tags, proximal and middle tags positively responding to stimulation are more than those negatively responding to stimulation (Supplemental Fig.3SC1). This result suggests that stimulation indeed raised transcription of tags but did not significantly trigger transcription switch between proximal and middle poly(A) sites. However, Supplemental Figures S3A2 and Supplemental Fig.3SA3 definitely show that stimulation remarkably increased transcription switch from distal poly(A) sites back to middle poly(A) site (Supplemental Fig.3SB2) or back to proximal poly(A) sites (Supplemental Fig.3SB3) while positively and negatively accordant transcriptions between proximal and distal poly(A) sites and between middle and distal poly(A) sites became very weak. In genes with proximal and middle poly(A) sites, positive tags were more than negative tags but in genes with distal poly(A) sites, negative tags were more than positive tags(Supplemental Fig.3SC2-3). These results strongly indicate that, as seen in genes with two tags, in genes with three tags, CD3/28cosimulation resulted in more usage of shorter tags.

## Discussion

Transcriptional profile via next generation sequencing technologies has underscored the complexity of transcript isoform variation in mammalian cells. The extent of this variation, whether the identity and function of a protein product (e.g. alternative splicing) or the visibility of the gene product is altered due to the posttranscriptional regulatory machinery (e.g. ACP), has been broadly appreciated. It is of great interest to leverage newer technologies to globally assess the impact of these processes on human disease, particularly since several individual examples in which dysregulation of either process contributes to human disease exist (Danckwardt et al. 2008).

The identification of relative expression changes of individual mRNA isoforms derived from a single transcript unit can in theory be performed by considering each of these isoforms from the transcript unit as a single entity and applying statistical approaches DESeq(Anders and Huber 2010), baySeq(Hardcastle and Kelly 2010), edgeR exact test (Robinson et al. 2009; Robinson and Smyth 2008), edgeR GLM (McCarthy et al. 2012), DEXSeq (Anders et al. 2012), Cuffdiff (Trapnell et al. 2013; Trapnell et al. 2010), DiffSplice(Hu et al. 2012), and SplicingCompass (Aschoff et al. 2013) to determine whether a given isoform is differentially expressed between two different cell states. However, these approaches do not directly assess whether isoforms within genes are associated with the conditions in transcription. Valid statistical approaches for doing so are Pearson chi-square test and Fisher exact test, but these methodologies cannot currently use information of variation in replicate libraries. As seen in Supplemental Appendix B, the Pearson chi-square of mean count over all given replicates is biased because mean noise cannot be excluded. Although Cochran Mantel Haenszel (CMH) chi-square test approach can be applied to repeat two-by-two table data or stratified count data and indeed its power is significantly higher than Pearson chi-square tests and Fisher Exact tests (see Supplemental Table 2) but our simulation studies show that its estimated FDR profile is still much higher than true FDR profile (see Supplemental Table 2 and Fig. S2).

Therefore, we need to develop a novel chi-square test methodology. Within the context of NGS datasets where individual transcript units have a potential to be characterized by the production of multiple mRNA isoforms, it is not clear what the relationship is between replication and the p-values of chi-square statistics. To avoid this puzzle issue, non-parametric approach is preferably chosen. This approach requires that an observed Pearson chi-square profile be compared to the null Pearson chi-square profile given a threshold. Since null chi-square profile does not exist, it must be estimated. In our methodology, we adopt a manner similar to estimation of null variance from replicate within-cell data to estimate the null chi-square profile. To choose appropriate threshold, we need to estimate FDR cutoff within which the findings declared by comparison between observed and null chi-square profiles are believed to be reliable.

While there are several alternative approaches for estimation of FDR, for example, the Benjamini-Hochberg procedure (Benjamini and Hochberg 1995; Benjamini and Yekutieli 2001), the Benjamini-Liu procedure (Benjamini and Liu 1999), and the Pounds–Cheng procedure (Pounds and Cheng 2006), Tan-Xu multi-procedures (Tan and Xu 2014), these FDR estimators are based on p-value profile and are not suitable to our non-parametric method. The permutation-based estimator developed by Tusher et al. (Tusher et al. 2001) has been shown both in theory and in simulation to be handicapped by bias in small sample sizes (Tan et al. 2006; Tan et al. 2008). Storey and Tibshriani (Storey and Tibshirani 2003) proposed a q-value as a new FDR estimator. Similarly, the q-value approach is also based on p-values (Dabney and Storey 2006; Storey and Tibshirani 2003). The other ranking analysis methods (Tan 2011; Tan et al. 2006; Tan et al. 2008) are designed to be applicable to continues data sets or cannot otherwise be used to this context. To accurately estimate FDR for the chi-square findings, we here proposed a novel approach. Our approach is based on a simple principle: given a threshold, tags were declared to be associated with conditions by comparing treatment ranking chi-square profile to null ranking chi-square profile and number of false tags associated with conditions was determined by comparing null chi-square at low ranking position to that at higher ranking position. For a given threshold Δ, a tag at *t*^*^ = *b*_Δ_ + *k* in a linear space of the null chi-squares has growth of probability to be declared as a false positive with increment of *k*. As seen in Figure 4, if a treated chi-square distribution falls into the null chi-square distribution, then we cannot find *a*_Δ_ value except for *a*_Δ_ = 1 across a set of given thresholds, and hence we also cannot find *b*_Δ_ value except for *b*_Δ_ = *S*. Thus, *N*_Δ_ = 0, *F*_Δ_ = 0, and FDR cannot be defined in such a case. Supplemental Figure S1 shows that estimated FDR is larger than its true FDR when threshold is small but it tends to be very close to its true value as threshold increases. This property guarantees that estimation of FDR is conservative and reliable at any threshold level.

## Materials and Methods

### Cell Lines and Stimulation

Human primary CD4 T cells were obtained from buffy coats derived from the Gulf Coast Regional Blood Bank via the EasySep Negative Selection Kit (Stem Cell Technologies) as per manufacturer’s instructions. Purity (>90%) was assessed via flow cytometry for CD3 and CD4 (UCHT1 and RPA-T4, respectively). Cells were maintained in RPMI (ATCC) with 10% fetal bovine serum supplemented with 10mM HEPES pH 7.4, non-essential amino acids, 2mM L-glutamate, 50 μM Beta-mercaptoethanol, 100 units/mL penicillin and 100 μg/mL streptomycin (all from Gibco/Life Technologies). T-cells were stimulated with plate-bound antibodies (1 μg/mL anti-CD3 (OKT3 – eBiosciences), 5 μg/mL anti-CD28 (CD28.2 – BD Pharmingen). Activation of T-lymphocytes was verified via flow cytometric detection of CD69 expression (FN-50) 16 hours after stimulation, and cells were harvested at 48 hours.

### High-throughput sequencing library generation

Total RNA was harvested from resting and stimulated cells with Trizol reagent (Life Technologies) as per manufacturer instructions. Polyadenylated RNA was isolated with the Poly(A)-Purist MAG (Ambion/Life Technologies) kit as per manufacturer instructions. High-throughput sequencing libraries were generated essentially as described (Shepard et al. 2011), with the exception that “barcoded” linkers were used to facilitate multiplexing. Libraries were sequenced via 50 bp paired-end sequencing on an Illumina GAIIx.

### Library processing and mapping

Paired end reads were mapped to the mm9 build of the mouse genome using the paired-end mapping module of bwa 0.5.9 (Li and Durbin 2009), default alignment stringency, and requiring that each read be mapped in a proper pair. To rescue reads crossing splicing junctions, non-mapping reads were remapped to the UCSC KnownGene reference and projected back to the mm9. Individual reads were condensed to tags based on their 3' coordinate using a sliding window of 20 nucleotides, using the frequency-weighted median 3' coordinate as the tag identifier. Tags were then filtered for false priming using a progressive filtering strategy assessing adenosine and guanine composition in the five, ten, and fifteen bases followed the tag-mapping site. Tags were assigned to individual transcription units based on UCSC KnownGene annotations. For each transcription unit, the aggregate tags mapping to the unit were ranked based on frequency. Tags were extracted from highest to lowest frequency until the extracted tags represented greater than 90% of the aggregate frequency for the gene. The remaining tags were discarded. Libraries were normalized using a negative binomial model within the DEseq package (Anders and Huber 2010). For “gene” analysis, the summed frequency of all tags mapping to each transcription unit was considered as a single entity.

### RAX2 Packages and statistical analysis

The methods described in this paper, including estimations of null chi-square distribution and FDR, are implemented in software package RAX2. RAX2 is written in R. The current version of RAX2 is designed to analyze count data in two distinct states and the results are output in MS DOS csv format. Performance of RAX2 and the other statistical analysis process are given in Supplemental Note S4. RAX2 package can be found in Supplemental Materials.

## Acknowledgements

We are indebted to Natee Kongchan and Arindam Chaudhury for library preparation and Alexander Ruch for pipeline construction. This work was funded by The Cancer Prevention and Research Institute of Texas (HIRP100475) and the National Cancer Institute (CA126752, CA131474).

## References

Anders, S. and Huber, W. 2010. Differential expression analysis for sequence count data. Genome Biol 11: R106.

Anders, S., Reyes, A., and Huber, W. 2012. Detecting differential usage of exons from RNA-seq data. Genome Res 22: 2008–2017.

Aschoff, M., Hotz-Wagenblatt, A., Glatting, K.H., Fischer, M., Eils, R., and Konig, R. 2013. SplicingCompass: differential splicing detection using RNA-seq data. Bioinformatics 29: 1141–1148.

Baggerly, K.A., Deng, L., Morris, J.S., and Aldaz, C.M. 2003. Differential expression in SAGE accounting for normal between-library variation. Bioinformatics 19: 1477–1483.

Benjamini, Y. and Hochberg, Y. 1995. Controlling the false discovery rate: a practical and powerful approach to multiple testing. J Roy Statist Soc Ser B (Methodological) 57: 289–300.

Benjamini, Y. and Liu, W. 1999. A step-down multiple hypotheses testing procedures that controls the false discovery rate under independence. Statist. Plann. Inference 82163–170.

Benjamini, Y. and Yekutieli, D. 2001. The control of the false discovery rate in multiple testing under dependency. Ann. Statist 29: 1165–1188.

Colgan, D.F. and Manley, J.L. 1997. Mechanism and regulation of mRNA polyadenylation. Genes Dev 11: 2755–2766.

Dabney, A.R. and Storey, J.D. 2006. A reanalysis of a published Affymetrix GeneChip control dataset. Genome Biol 7: 401.

Danckwardt, S., Hentze, M.W., and Kulozik, A.E. 2008. 3' end mRNA processing: molecular mechanisms and implications for health and disease. EMBO J 27: 482–498.

Flavell, S.W., Kim, T.K., Gray, J.M., Harmin, D.A., Hemberg, M., Hong, E.J., Markenscoff-Papadimitriou, E., Bear, D.M., and Greenberg, M.E. 2008. Genome-wide analysis of MEF2 transcriptional program reveals synaptic target genes and neuronal activity-dependent polyadenylation site selection. Neuron 60: 1022–1038.

Hardcastle, T.J. and Kelly, K.A. 2010. baySeq: empirical Bayesian methods for identifying differential expression in sequence count data. BMC Bioinformatics 11: 422.

Hu, Y., Huang, Y., Du, Y., Orellana, C.F., Singh, D., Johnson, A.R., Monroy, A., Kuan, P.F., Hammond, S.M., Makowski, L. et al. 2012. DiffSplice: the genome-wide detection of differential splicing events with RNA-seq. Nucleic Acids Res 41: e39.

Hunt, A.G., Xu, R., Addepalli, B., Rao, S., Forbes, K.P., Meeks, L.R., Xing, D., Mo, M., Zhao, H., Bandyopadhyay, A. et al. 2008. Arabidopsis mRNA polyadenylation machinery: comprehensive analysis of protein-protein interactions and gene expression profiling. BMC Genomics 9: 220.

Ji, Z., Lee, J.Y., Pan, Z., Jiang, B., and Tian, B. 2009. Progressive lengthening of 3' untranslated regions of mRNAs by alternative polyadenylation during mouse embryonic development. Proc Natl Acad Sci U S A 106: 7028–7033.

Lee, J.Y., Yeh, I., Park, J.Y., and Tian, B. 2007. PolyA_DB 2: mRNA polyadenylation sites in vertebrate genes. Nucleic Acids Res 35: D165–168.

Li, H. and Durbin, R. 2009. Fast and accurate short read alignment with Burrows-Wheeler transform. Bioinformatics 25: 1754–1760.

MacDonald, C.C. and McMahon, K.W. 2011. Tissue-specific mechanisms of alternative polyadenylation: testis, brain, and beyond. Wiley Interdiscip Rev RNA 1: 494–501.

Mayr, C. and Bartel, D.P. 2009. Widespread shortening of 3'UTRs by alternative cleavage and polyadenylation activates oncogenes in cancer cells. Cell 138: 673–684.

McCarthy, D.J., Chen, Y., and Smyth, G.K. 2012. Differential expression analysis of multifactor RNA-Seq experiments with respect to biological variation. Nucleic Acids Res 40: 4288–4297.

Meyers, B.C., Vu, T.H., Tej, S.S., Ghazal, H., Matvienko, M., Agrawal, V., Ning, J., and Haudenschild, C.D. 2004. Analysis of the transcriptional complexity of Arabidopsis thaliana by massively parallel signature sequencing. Nat Biotechnol 22: 1006–1011.

Pounds, S. and Cheng, C. 2006. Robust estimation of the false discovery rate. Bioinformatics 1979–1987.

Robinson, M.D., McCarthy, D.J., and Smyth, G.K. 2009. edgeR: a Bioconductor package for differential expression analysis of digital gene expression data. Bioinformatics 26: 139–140.

Robinson, M.D. and Smyth, G.K. 2008. Small-sample estimation of negative binomial dispersion, with applications to SAGE data. Biostatistics 9: 321–332.

Sandberg, R., Neilson, J.R., Sarma, A., Sharp, P.A., and Burge, C.B. 2008. Proliferating cells express mRNAs with shortened 3' untranslated regions and fewer microRNA target sites. Science 320: 1643–1647.

Shen, Y., Ji, G., Haas, B.J., Wu, X., Zheng, J., Reese, G.J., and Li, Q.Q. 2008. Genome level analysis of rice mRNA 3'-end processing signals and alternative polyadenylation. Nucleic Acids Res 36: 3150–3161.

Shen, Y., Venu, R.C., Nobuta, K., Wu, X., Notibala, V., Demirci, C., Meyers, B.C., Wang, G.L., Ji, G., and Li, Q.Q. 2011. Transcriptome dynamics through alternative polyadenylation in developmental and environmental responses in plants revealed by deep sequencing. Genome Res 21: 1478–1486.

Shepard, P.J., Choi, E.A., Lu, J., Flanagan, L.A., Hertel, K.J., and Shi, Y. 2011. Complex and dynamic landscape of RNA polyadenylation revealed by PAS-Seq. RNA 17: 761–772.

Shi, Y., Di Giammartino, D.C., Taylor, D., Sarkeshik, A., Rice, W.J., YatesJ.R., 3rd, Frank, J., and Manley, J.L. 2009. Molecular architecture of the human pre-mRNA 3' processing complex. Mol Cell 33: 365–376.

Storey, J.D. and Tibshirani, R. 2003. Statistical significance for genomewide studies. Proc Natl Acad Sci U S A 100: 9440–9445.

Tan, Y.D. 2011. Ranking analysis of correlation coefficients in gene expressions. Genomics 97:58–68.

Tan, Y.D., Fornage, M., and Fu, Y.X. 2006. Ranking analysis of microarray data: a powerful method for identifying differentially expressed genes. Genomics 88: 846–854.

Tan, Y.D., Fornage, M., and Xu, H. 2008. Ranking analysis of F-statistics for microarray data. BMC Bioinformatics 9: 142.

Tan, Y.D. and Xu, H. 2014. A general method for accurate estimation of false discovery rates in identification of differentially expressed genes. Bioinformatics 30: 2018–2025.

Tian, B., Pan, Z., and Lee, J.Y. 2007. Widespread mRNA polyadenylation events in introns indicate dynamic interplay between polyadenylation and splicing. Genome Res 17: 156–165.

Trapnell, C., Hendrickson, D.G., Sauvageau, M., Goff, L., Rinn, J.L., and Pachter, L. 2013. Differential analysis of gene regulation at transcript resolution with RNA-seq. Nat Biotechnol 31: 46–53.

Trapnell, C., Williams, B.A., Pertea, G., Mortazavi, A., Kwan, G., van Baren, M.J., Salzberg, S.L., Wold, B.J., and Pachter, L. 2010. Transcript assembly and quantification by RNA-Seq reveals unannotated transcripts and isoform switching during cell differentiation. Nat Biotechnol 28: 511–515.

Tusher, V.G., Tibshirani, R., and Chu, G. 2001. Significance analysis of microarrays applied to the ionizing radiation response. Proc Natl Acad Sci U S A 98: 5116–5121.

Wang, E.T., Sandberg, R., Luo, S., Khrebtukova, I., Zhang, L., Mayr, C., Kingsmore, S.F., Schroth, G.P., and Burge, C.B. 2008. Alternative isoform regulation in human tissue transcriptomes. Nature 456: 470–476.

Winter, J., Kunath, M., Roepcke, S., Krause, S., Schneider, R., and Schweiger, S. 2007. Alternative polyadenylation signals and promoters act in concert to control tissue-specific expression of the Opitz Syndrome gene MID1. BMC Mol Biol 8: 105.

Wu, X., Liu, M., Downie, B., Liang, C., Ji, G., Li, Q.Q., and Hunt, A.G. 2011. Genome-wide landscape of polyadenylation in Arabidopsis provides evidence for extensive alternative polyadenylation. Proc Natl Acad Sci U S A 108: 12533–12538.

Yu, M., Sha, H., Gao, Y., Zeng, H., Zhu, M., and Gao, X. 2006. Alternative 3' UTR polyadenylation of Bzw1 transcripts display differential translation efficiency and tissue-specific expression. Biochem Biophys Res Commun 345: 479–485.

Zhao, J., Hyman, L., and Moore, C. 1999. Formation of mRNA 3' ends in eukaryotes: mechanism, regulation, and interrelationships with other steps in mRNA synthesis. Microbiol Mol Biol Rev 63: 405–445.

